# Comprehensive biological interpretation of gene signatures using semantic distributed representation

**DOI:** 10.1101/846691

**Authors:** Yuumi Okuzono, Takashi Hoshino

## Abstract

Recent rise of microarray and next-generation sequencing in genome-related fields has simplified obtaining gene expression data at whole gene level, and biological interpretation of gene signatures related to life phenomena and diseases has become very important. However, the conventional method is numerical comparison of gene signature, pathway, and gene ontology (GO) overlap and distribution bias, and it is not possible to compare the specificity and importance of genes contained in gene signatures as humans do.

This study proposes the gene signature vector (GsVec), a unique method for interpreting gene signatures that clarifies the semantic relationship between gene signatures by incorporating a method of distributed document representation from natural language processing (NLP). In proposed algorithm, a gene-topic vector is created by multiplying the feature vector based on the gene’s distributed representation by the probability of the gene signature topic and the low frequency of occurrence of the corresponding gene in all gene signatures. These vectors are concatenated for genes included in each gene signature to create a signature vector. The degrees of similarity between signature vectors are obtained from the cosine distances, and the levels of relevance between gene signatures are quantified.

Using the above algorithm, GsVec learned approximately 5,000 types of canonical pathway and GO biological process gene signatures published in the Molecular Signatures Database (MSigDB). Then, validation of the pathway database BioCarta with known biological significance and validation using actual gene expression data (differentially expressed genes) were performed, and both were able to obtain biologically valid results. In addition, the results compared with the pathway enrichment analysis in Fisher’s exact test used in the conventional method resulted in equivalent or more biologically valid signatures. Furthermore, although NLP is generally developed in Python, GsVec can execute the entire process in only the R language, the main language of bioinformatics.

## Introduction

The recent rise of microarray and next-generation sequencing (NGS) in genome-related fields has made it possible to easily acquire gene expression data at the whole gene level. As a result, interpretation of life phenomena and diseases is advancing [1].

To identify the gene population involved in a phenotype, gene expression data for comparison between healthy subjects and subjects with diseases as well as treated and untreated groups can be obtained. Based on the correlation between the representative expression value of the gene signature and the phenotype, the gene signature of genes related to the phenotype can be identified, and the biological interpretation of gene signatures can be performed.

To interpret a gene signature identified in this data-driven manner, it is necessary to avoid bias due to the large number of genes that must be interpreted and comprehensiveness and completeness of human knowledge. Therefore, interpretation is commonly performed by comparing the gene signature, such as differentially expressed genes and gene modules, against a biological gene signature database (such as pathway and GO) and identifying an objective association from a biological perspective [2].

Numerous methodologies for association with pathways have been proposed. Common examples include Fisher’s exact test, which is a classical statistical test for the specific overlap of genes; over-representation analysis and gene set enrichment analysis [3], which statistically process the number of overlapping genes and ranking bias by incorporating randomization; and modular enrichment analysis and EnrichNet with graph-based statistics of biological networks [4, 5].

However, these comparisons are numerical, and it is thus not possible to compare the semantic nuances of the included genes as humans do. After performing the aforementioned analyses, to interpret the gene signature it is necessary to perform a number of comprehensive judgments to identify whether the genes overlapped by humans are specific to the pathway, determine their biological importance, and establish whether they contain genes that have similar meaning without direct overlap.

In this study, we propose a novel method for associating biological gene signatures by applying a distributed representation model [6] of documents to facilitate the interpretation of gene signatures (see Fig 1). The distributed representation of documents is a technology for vectorizing documents of any length that was developed in the field of natural language processing (NLP). Words (that are included in documents) and documents (that are sets of words) are vectorized to convert a semantic expression of the words and documents into a mathematical expression that can be easily processed by a computer. As the first step, feature extraction is performed at the word level, and as the second step, feature extraction is performed at the document level as a set of words. Because feature vectors can be semantically compared by performing mathematical comparisons, they are used in a variety of real-world situations, such as sentence classification, content recommendation, sentiment analysis, and spam filtering [7].

**Fig 1.**
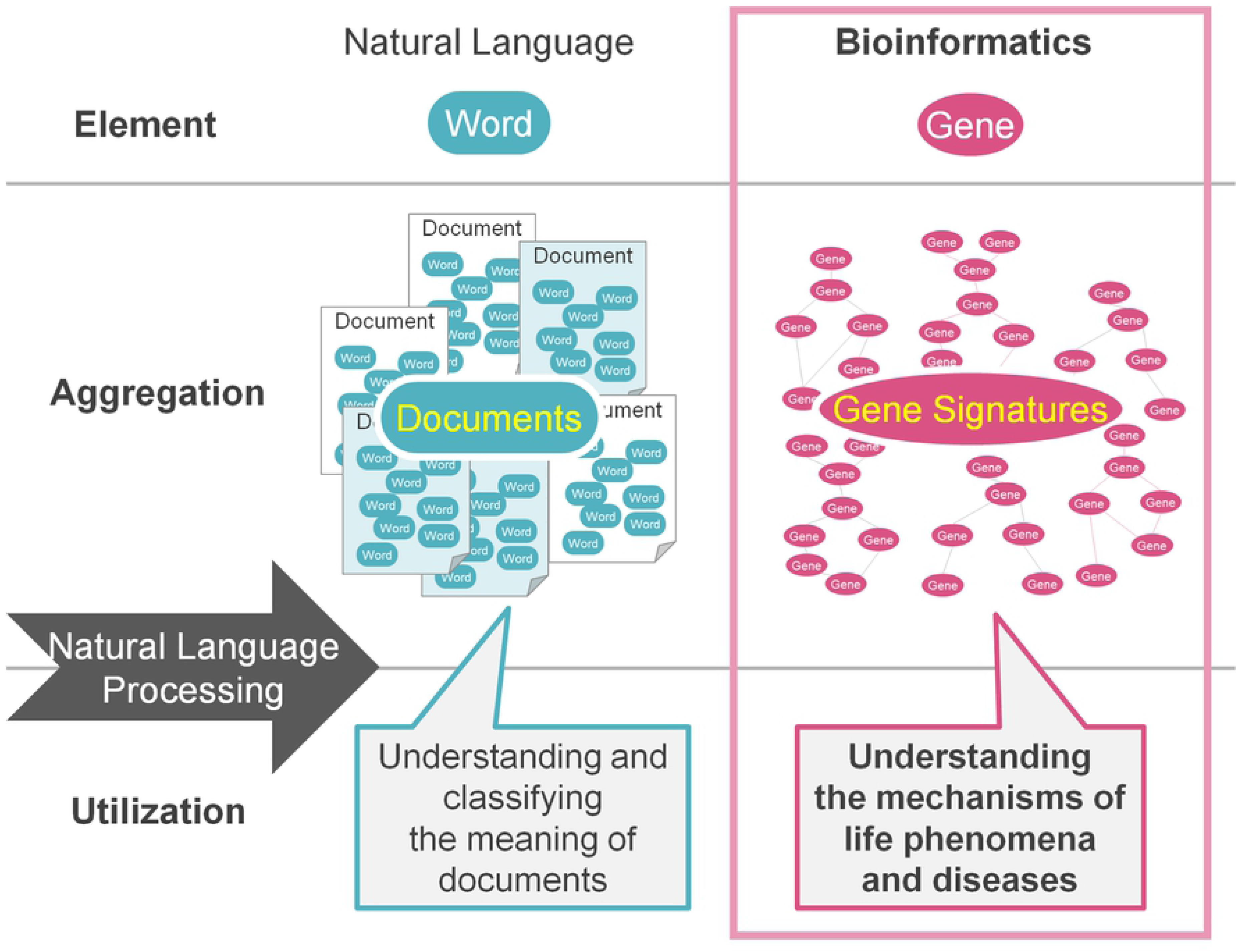
Research concept.

Beginning with Doc2Vec [8], which used a distributed representation of words, innovative techniques related to the distributed expression of a large number of sentences have been proposed in the past several years, and the accuracy of document interpretation has improved [9]. Typical methods of distributed representation of documents include statistical semantic extraction methods [10], methods that combine distributed representations of words [11] into document representations [12], methods that directly compress word and document IDs [8], methods of summing word vectors by multiplying the topics and specificities in the documents [9].

There are other NLP methods for the distributed representation of documents; however, the methods applicable to bioinformatics are limited, as there are differences in assumptions between NLP and bioinformatics. A meaningful component in bioinformatics is a gene, which corresponds to a word in NLP. In addition, a gene signature, which is a set of genes, corresponds to a document, which is a set of words. However, in NLP, the order and context of words are important, whereas the order of gene signatures compiled in a general pathway database often has no meaning.

In this study, we developed an original method for the interpretation of gene signatures by applying a distributed expression algorithm. The algorithm extracts semantic features of genes and their biological gene signatures and reveals specific relationships by comparing the abovementioned signatures with the gene signatures to be interpreted (e.g., differentially expressed genes, gene modules). As training data, a gene signature was used, which has a clear biological meaning in the molecular signatures database (MSigDB) [13] used in the conventional pathway enrichment analysis. Furthermore, Python is the primary programming language used for machine learning and NLP analysis; however, our proposed method can execute the entire process in R, which is the primary language in the analysis domain of bioinformatics. Therefore, the proposed method can be immediately used without further modification for bioinformatics analysis. Combining our proposed method of biological gene signature vectorization with conventional enrichment analysis can allow for more intuitive and reliable interpretation of gene signatures.

## Results

### Construction of gene and gene signature feature vectors by distributed representation

In this study, we developed a method for creating gene signature feature vectors and clarifying semantic similarity by applying methodology from the field of NLP (Fig 2). We defined proprietary functions using the packages published on the Comprehensive R Archive Network that can be used in the R language. In addition, we executed an original algorithm for creating a unique gene signature feature vector based on the sparse composite document vectors (SCDV) [9] method from NLP using only R language operations.

**Fig 2.**
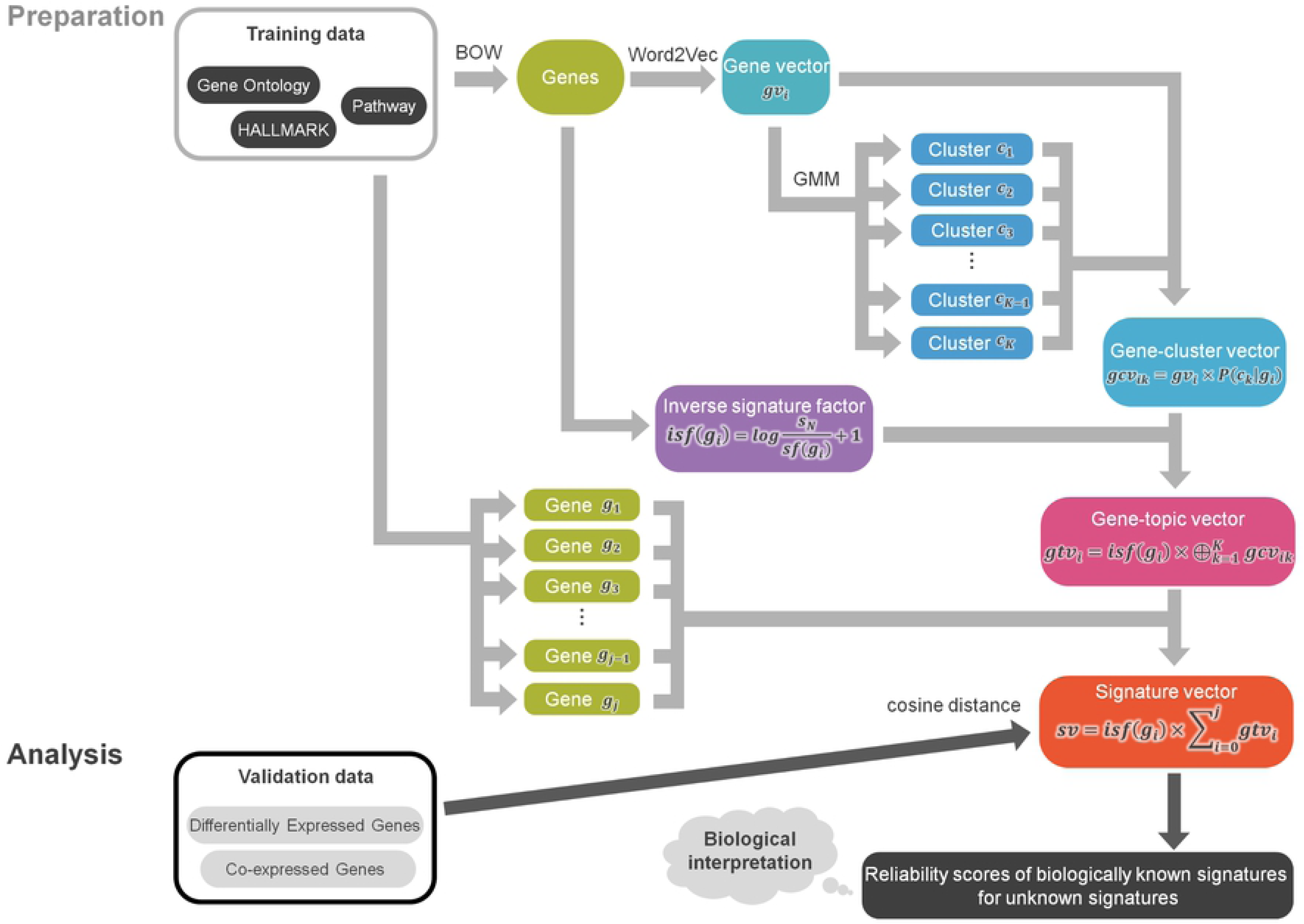
Algorithm and workflow of GsVec. GsVec is divided into preparation and analysis parts. In the preparation part, gene vectors are created from training data with a clear biological meaning (e.g., GO, Pathway, and Hallmark genes) using BOW and Word2Vec, and the probability of each cluster calculated by GMM is multiplied. Furthermore, gene-topic vectors are created by multiplying the inverse signature factor and averaging for each gene included in the gene signature. In the analysis part, validation data, which are not biologically interpreted gene signatures (e.g., differentially expressed genes and gene modules) are converted into signature vectors from the gene-topic vector created using training data. The cosine similarity between the validation and training data is calculated to obtain the association with biological meaning. *gv_i_* represents the vector of an arbitrary gene *g_i_*, *c* represents the cluster, *K* represents the number of clusters, *S_N_* represents the total number of gene signatures, *Sf* (*g_i_*) represents the number of gene signatures including gene *g_i_*, and ⊕ represents the concatenation.

The training data used 5,456 gene signatures (i.e., C2: Canonical pathway and C5: GO biological process), which were used in conventional pathway/GO enrichment.

A gene × gene signature matrix was created for the gene signature (equivalent to the Bags of Word step in NLP), and gene features were expressed using a distributed word representation algorithm [14] to create gene vectors (equivalent to Word2Vec processing in NLP). The clusters corresponding to the topics present in these gene signatures were extracted by soft clustering of the gene vectors with a Gaussian mixture model (GMM) [15, 16]. The probabilities that each word contributes to each cluster were multiplied for each cluster to obtain the abovementioned gene vectors, and those vectors were combined for each gene to obtain gene-cluster vectors.

Simultaneously, by dividing the total number of gene signatures by the number of gene signatures for each gene that contain that gene, the scores for reducing the weight of the gene appearing in various signatures were calculated. Hereinafter, this is referred to as the inverse signature factor, which is equivalent to IDF in NLP [17]. Gene-topic vectors were obtained by multiplying the abovementioned gene-cluster vectors by those scores. The signature vectors were obtained by averaging the gene-topic vectors for the genes included in each individual gene signature. The signature vectors which were feature vectors in the genes and the abovementioned gene signatures were used as training data in the subsequent analysis.

### Evaluation of gene signature vector (GsVec) performance using gene signatures with known biological interpretation

In this study, 5,242 gene signatures, excluding gene signatures derived from the BioCarta database from the C2 canonical pathway and C5 GO biological process, were used as training data. BioCarta’s 214-gene signatures were used as validation data with a known biological meaning. Furthermore, in both the training and validation data, we selected gene signatures related to immune function by human selection and tagged them. The number of applicable gene signatures was 405.

Signature vectors of the validation data were created from the gene-topic vectors created during learning in the same way as the training data, and the degrees of relevance with the training data were evaluated by the cosine similarity score with the learning set (hereinafter, the result of the cosine similarity calculated by a series of operations is referred to as *GsVec*). First, the GsVec results (i.e., similarity relationship between signature vectors) were visualized by two-dimensionally projecting them using t-distributed Stochastic Neighbor Embedding (t-SNE) [18] (Fig 3[A]). There was a tendency for immune-related signatures to be consolidated in one location. The abovementioned tendency was observed not only in the training data, but also in the validation data, and the meaning of the validation data was correctly predicted by GsVec. These results demonstrate that although GsVec using NLP is an entirely different approach from conventional methods, it can identify groups with similar meanings.

**Fig 3.**
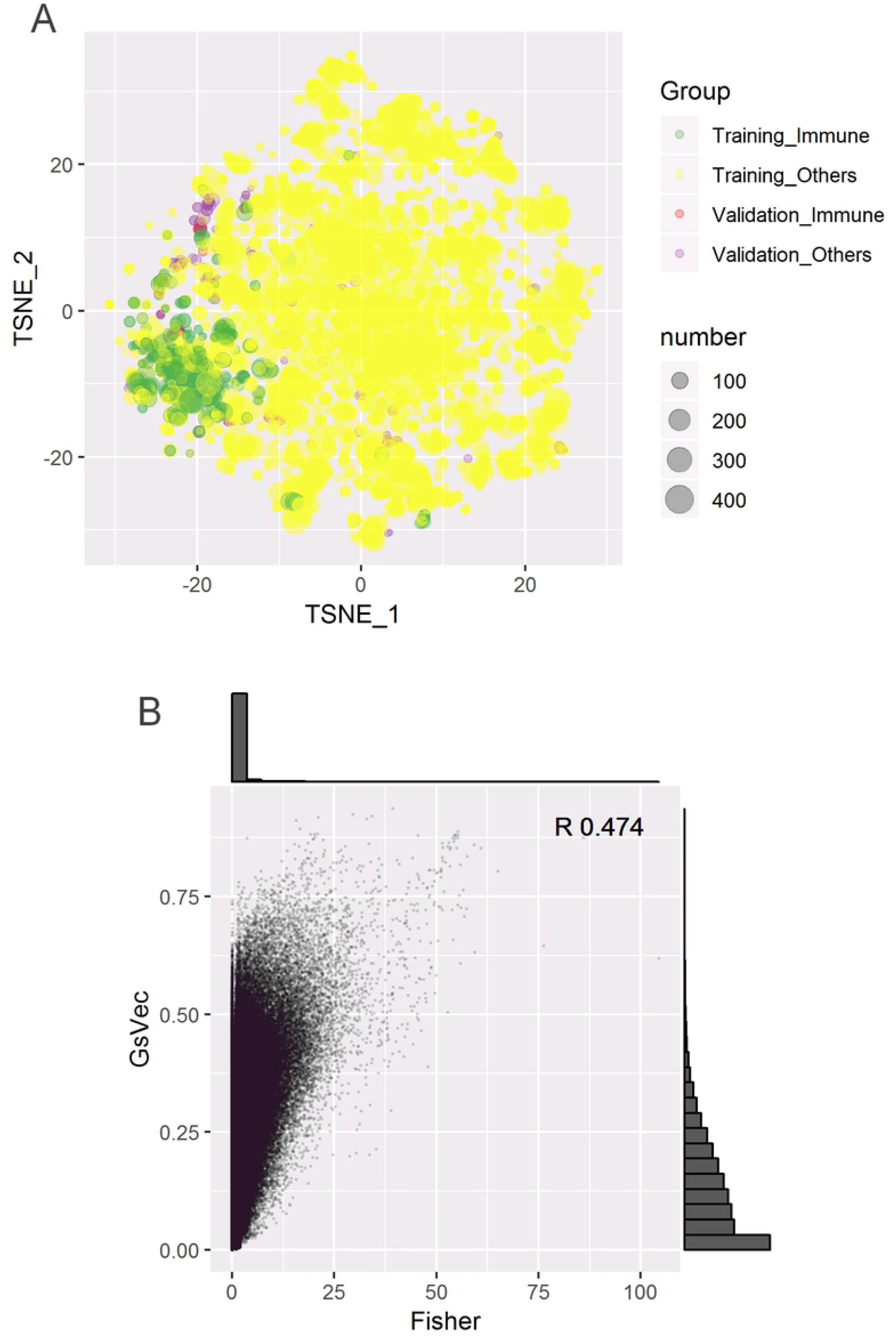
GsVec accuracy evaluation and comparison with Fisher. A) Matrix of the cosine distance between signature vectors of training and validation data projected in two dimensions with t-SNE. The size of the circle represents the number of genes included in each signature. Green color refers to training data related to immunity, purple to training data not related to immunity, blue to validation data related to immunity, and red to validation data not related to immunity. B) Scatter plot by the -log10 P-value of Fisher and the cosine distance between the signature vectors of the training and validation data. A histogram of the distribution of each value is also illustrated. The R in the upper right of the scatter plot indicates the Pearson correlation coefficient between GsVec and Fisher.

Next, the P-value of the overlap between Fisher’s exact test training data and validation data was converted to a -log10 value (hereinafter, −log10 (P-value) obtained by Fisher’s exact test is referred to as *Fisher*) and compared with the GsVec results. The correlation by Pearson coefficient score between GsVec and Fisher was 0.453, and the relationship with a score of 0.75 or higher in GsVec was a significant score less than 0.001 (-log10 (4)) in Fisher. In addition, the degree of relevance was concentrated in the Fisher distribution near 0, which was not significant, whereas in GsVec, the distribution had a long tail. These results suggest that GsVec was able to reflect robust results that were significant in Fisher, and could also associate gene signatures with interpretation that were difficult to interpret in Fisher (Fig 3[B]).

In addition, the relationship between the gene signatures of individual validation data and training data was compared between GsVec and Fisher. The top 10 results were the same for GsVec and Fisher for gene signatures, with a large number of genes included in the gene signature in the training data and a large number of overlaps. Furthermore, in GsVec, even if a gene included in the gene signature in the training data was small and the number of overlapped genes was small, if the overlapped genes were characteristic genes in the signature vector space, there was a tendency to display high relevance. Typical results are presented in Fig 4.

**Fig 4.**
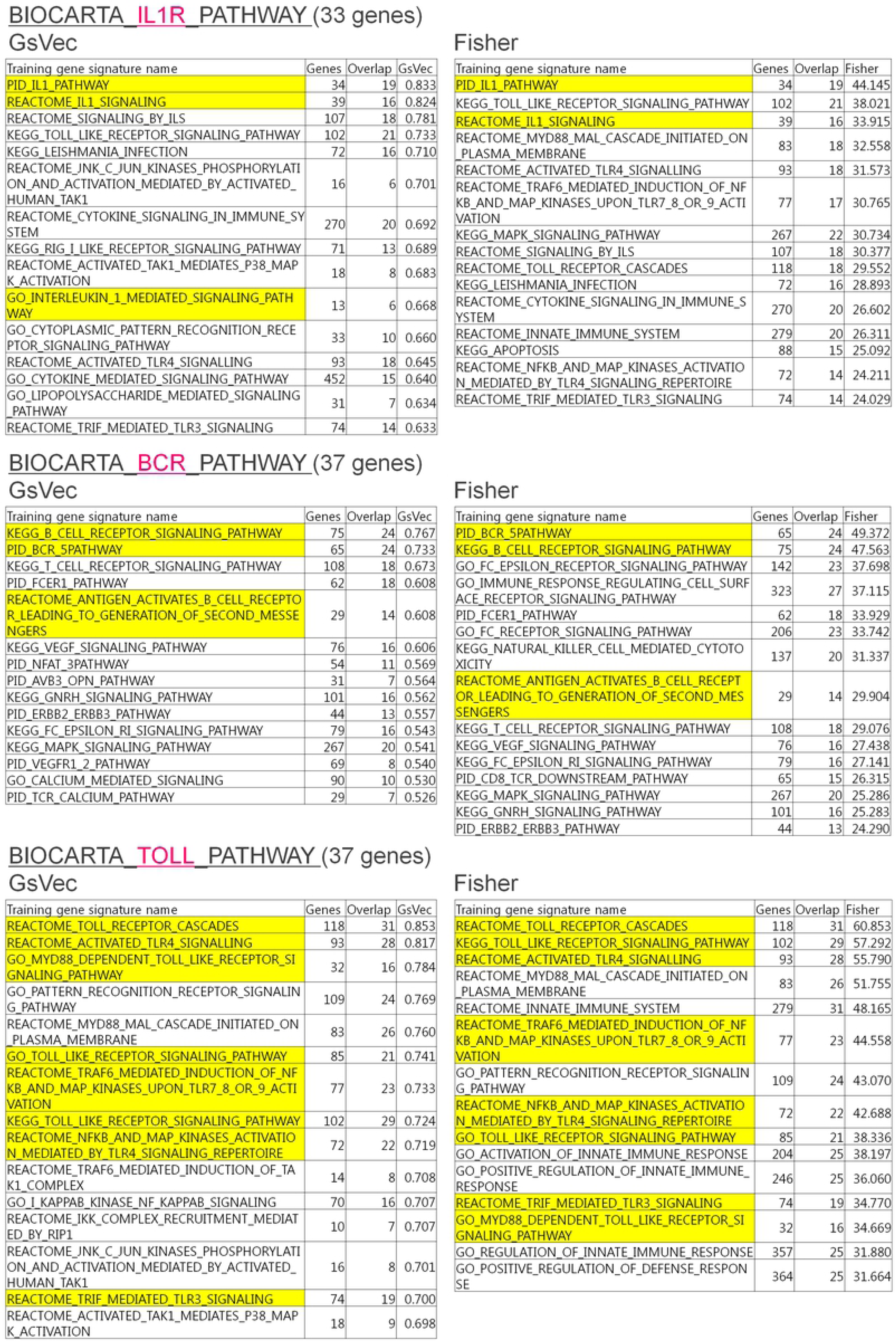
Comparison between GsVec and Fisher on top 15 results with high relevance between training data and validation data. Each top 15 result that was highly correlated with the gene signature and GsVec of the validation data or that was highly significant with Fisher is presented in the left and right tables. The column *Training gene signature name* indicates the name of the corresponding training data, and the biological data that directly match the validation data are highlighted in yellow. The *genes* column contains the total number of genes included in the gene signature of the corresponding training data, the *overlap* column contains the number of genes overlapped between the corresponding validation data and training data, and the *Fisher’s* column represents the -log10 (P-value) of the significance of the corresponding training data and validation data by Fisher’s exact test. The *GsVec* column exhibits similarities according to the cosine distance of the signature vectors of the corresponding training and validation data.

Here, we examined whether GsVec was able to produce biologically meaningful insights using three representative examples of immune-related pathways. In the Interleukin-1 receptor (IL1R) pathway, the direct gene signature name *IL1* or *Interleukin 1* could be identified in both GsVec and Fisher. Similar results were observed in the B cell receptor and Toll pathways, including *Toll-like receptor*. In addition, gene signature names indicating broader concepts, such as *cytokine signaling* and *pattern recognition receptor*, were observed in the IL1R and Toll pathways, respectively. These were extracted by GsVec and are associated with biologically relevant gene signatures.

### Association of biological meaning with GsVec using differentially expressed genes (DEGs) from real data

Data-driven gene signatures, such as DEGs in the affected tissues and normal tissues of diseases published in the Expression Atlas [19] and DEGs extracted from The Cancer Genome Atlas (TCGA) [20] data, were analyzed with GsVec and Fisher (Fig 5). In all datasets, DEGs were used; they were upregulated in the affected tissue compared to the normal tissue. However, because in cancer there were too many DEGs (approximately 1,000– 2,000) for the pathway enrichment analysis, only the results of other diseases were interpreted. Additionally, only in multiple sclerosis (MS), the number of DEGs was low; thus, we analyzed not only upregulated (up) but also downregulated (down) DEGs. The individual analysis results are presented below.

**Fig 5.**
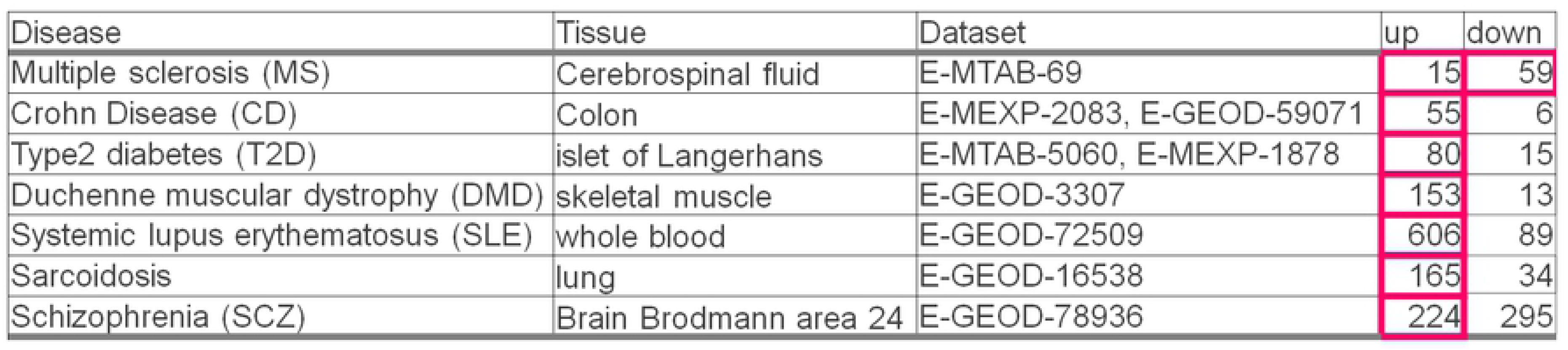
List of DEGs used for verification. The table illustrates the DEG data of affected tissues and control tissues of the major public diseases used as data-driven signatures, not signatures compiled in terms of biological meaning. The *up* column contains the number of DEGs that were higher than the control for the disease, while the *down* column contains the DEG numbers that were lower than the control for the disease. The red frames represent subsequent interpretations.

### Multiple sclerosis (MS) cerebrospinal fluid (CSF; E-MTAB-69)

In MS (up), B-cell-related signatures were identified in GsVec and Fisher (Fig 6). Although the involvement of T cells in the pathology of MS is well known, the participation of B cells has also attracted attention in recent years, and the possibility of therapeutic drugs targeting B cells has also been investigated [21, 22]. In contrast, in MS (down), the gene signature names *locomotion*, *taxis*, or *migration*, reminiscent of cell migration, were highly ranked. In this study, the dataset consisted of CSF-derived samples. The results indicated that the involvement of immune cells in the peripheral and central nervous system (CNS) was captured. Notably, the gene signature name, *nervous system development*, was highly ranked only in GsVec. In other words, when considering the pathological condition of MS, GsVec captured not only immunological but also neuronal aspects and was able to extract more biologically valid signatures.

**Fig 6.**
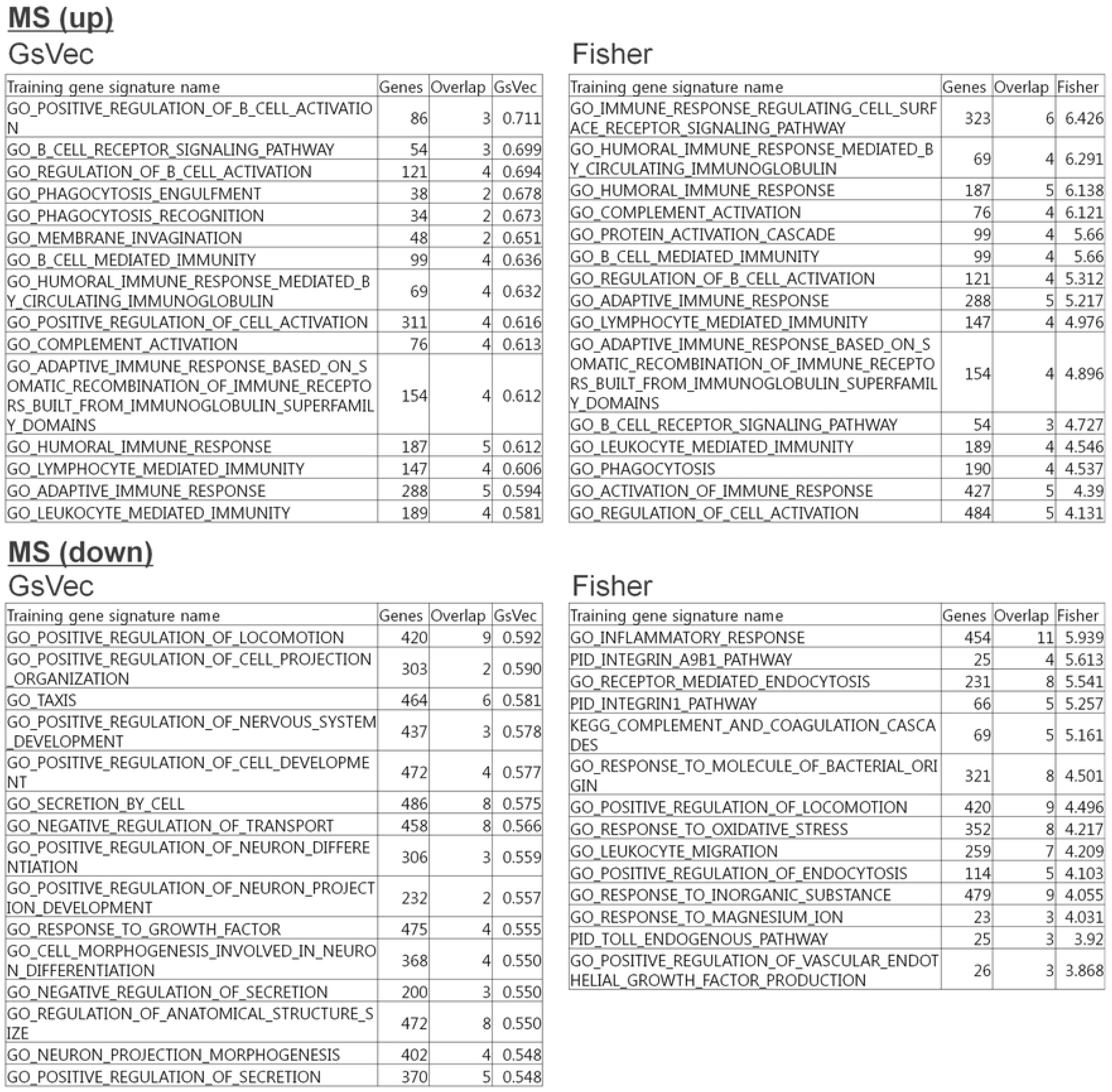
Comparison between GsVec and Fisher of top 15 results with high relevance in DEGs of multiple sclerosis (MS). Using the DEGs of MS cerebrospinal fluid (E-MTAB-69) as validation data, each top 15 result that was highly correlated with GsVec or highly significant with Fisher is presented in the left and right tables. The *genes* column contains the total number of genes included in the gene signature of the corresponding training data, the *overlap* column contains the number of genes overlapped between the corresponding validation and training data, and the *Fisher’s* column represents -log10 (P-value) of the significance of the corresponding training data and validation data by Fisher’s exact test. The *GsVec* column exhibits similarities according to the cosine distance of the signature vectors of the corresponding training and validation data.

### Crohn’s disease (CD) colon (E-MEXP-2083, E-GEOD-59071)

We examined a dataset of fresh frozen ileum mucosal tissue from CD patients, and gene signatures, such as *interferon gamma*, *cell adhesion*, and *leukocyte migration* were highly ranked in GsVec (see Fig 7). For inflammatory bowel diseases, such as CD and ulcerative colitis, it has been reported that intestinal immune cell trafficking has been identified as a central event in the pathogenesis of diseases. Additionally, cell adhesion is a pivotal step in several aspects of immune cell trafficking [23]. In Fisher, *interferon gamma* was highly ranked, and the broader concept *cytokine* was also highly ranked. However, the gene signature *asthma*, which may not be directly related to CD, was also highly ranked in Fisher. It has been reported that there are many common risk factors for the association between asthma and CD, including genetic and environmental factors [24]. Thus, GsVec and Fisher in CD displayed a similar trend but identified different characteristics on the whole. GsVec can extract more biologically meaningful signatures.

**Fig 7.**
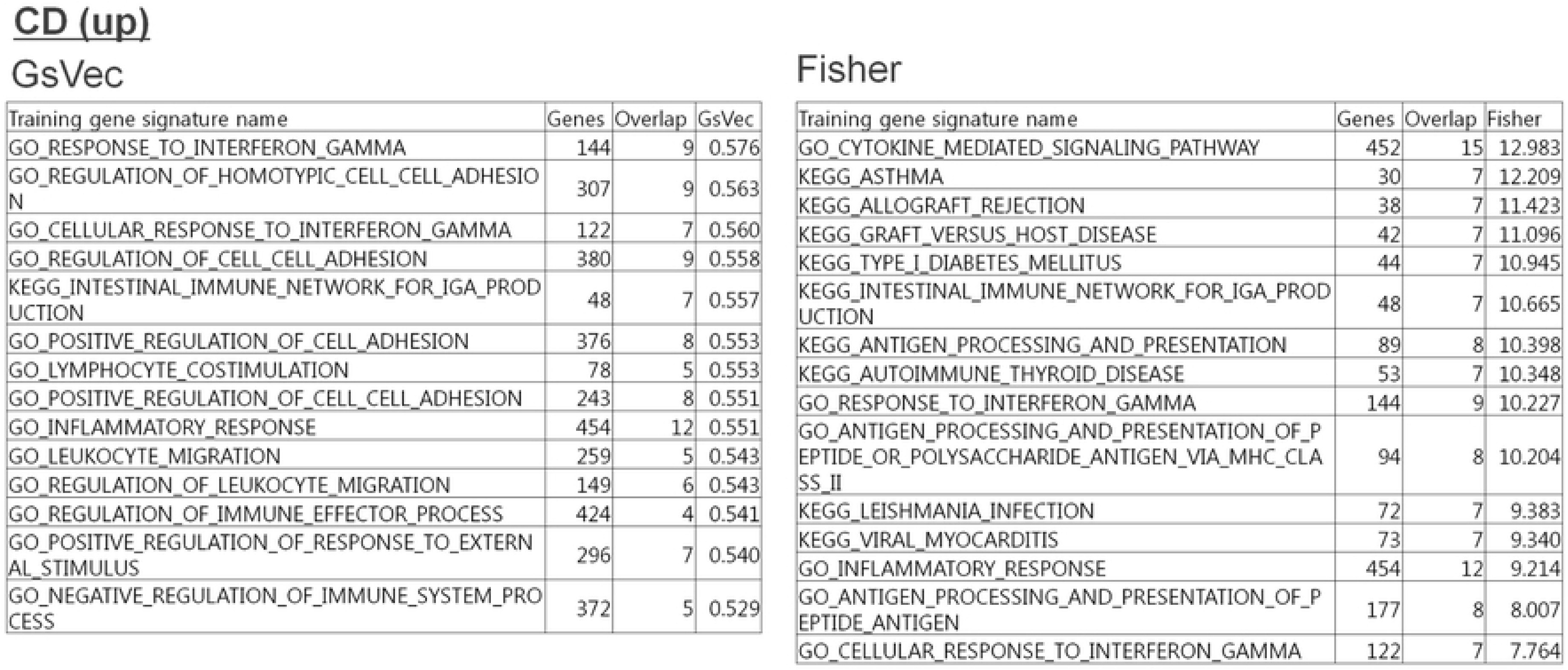
Comparison between GsVec and Fisher of top 15 results with high relevance in DEGs of Crohn’s disease (CD). Using the DEGs of CD colon (E-MEXP-2083, E-GEOD-59071) as validation data, each top 15 result that was highly correlated with GsVec or highly significant with Fisher is presented in the left and right tables. The details of each column are identical to those in Fig 6.

### Type 2 diabetes (T2D) islet of Langerhans (E-MTAB-5060)

In this study, we used a dataset consisting of islet of Langerhans tissue from healthy donors and T2D patients. In both GsVec and Fisher, gene signatures related to inflammation such as *inflammatory response*, *migration*, and *wound* were highly ranked and demonstrated a similar trend (see Fig 8). Insulin resistance and beta cell dysfunction are well known in T2D pathologies, and inflammation is related to the pathogenesis of these conditions [25]. In addition, injury and wound healing processes associated with the term *wounding* are known to alter responses to growth factors and cytokines in addition to tissue remodeling through cell migration and proliferation [26, 27]. These scientific reports examined the effect of macrophages on T2D-related ulcers and skin wounds, but not the islets themselves. The direct cause-and-effect relationship is unknown; however, based on the GO term definition, the relationship was linked to a gene signature involved in damaged tissue and tissue repair.

**Fig 8.**
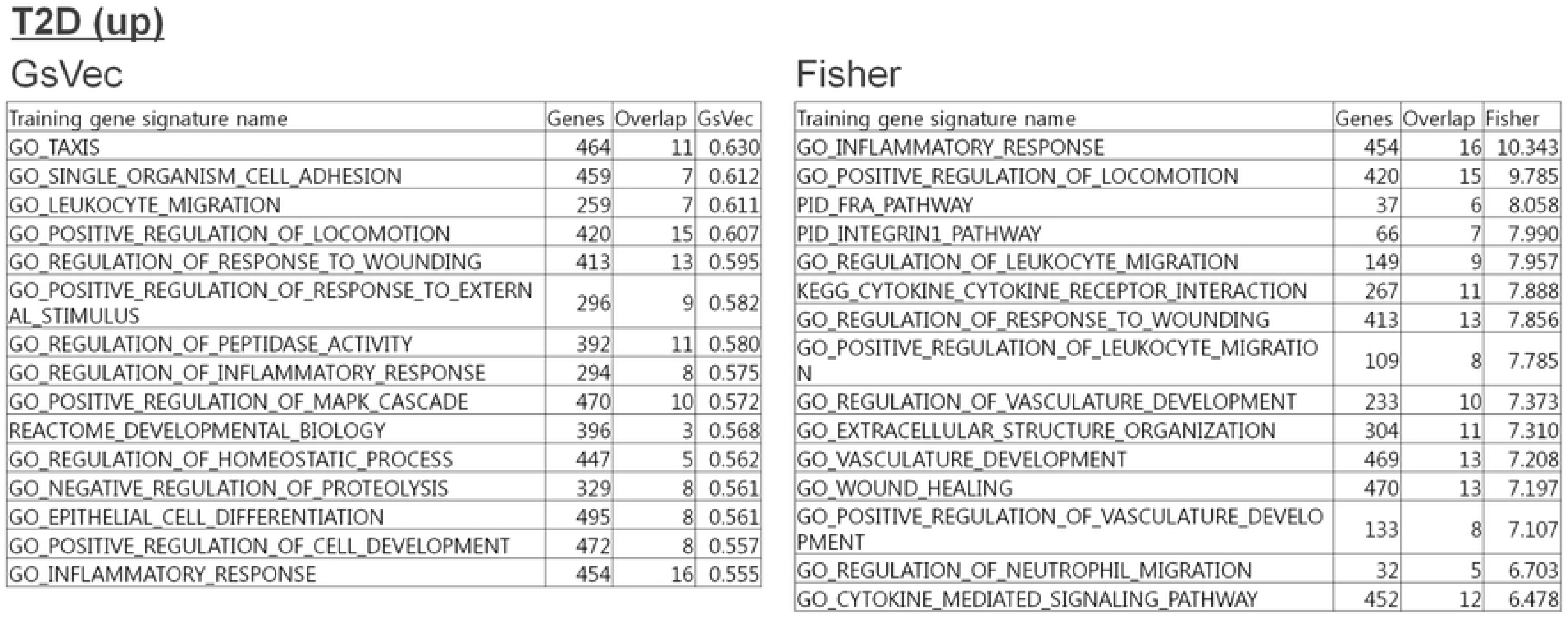
Comparison between GsVec and Fisher of top 15 results with high relevance in DEGs of Type 2 diabetes (T2D). Using the DEGs of T2D islet of Langerhans (E-MTAB-5060) as validation data, each top 15 result that had high correlation with GsVec or high significance with Fisher is presented in the left and right tables.

### Duchenne muscular dystrophy (DMD) skeletal muscle (E-GEOD-3307)

We then examined DMD, and the results demonstrated that GsVec and Fisher displayed similar trends (Fig 9). Specifically, gene signatures such as *extracellular structure organization* related to the extracellular matrix and *ossification* related to the bone were highly ranked. DMD is an inherited muscular disorder known to be caused by an abnormality in dystrophin, a cytoskeletal protein, and has been linked to extracellular matrix-related molecules [28]. In addition, a relationship between bone morphogenetic proteins signals and this disease has been reported, and several biological features have been extracted [29].

**Fig 9.**
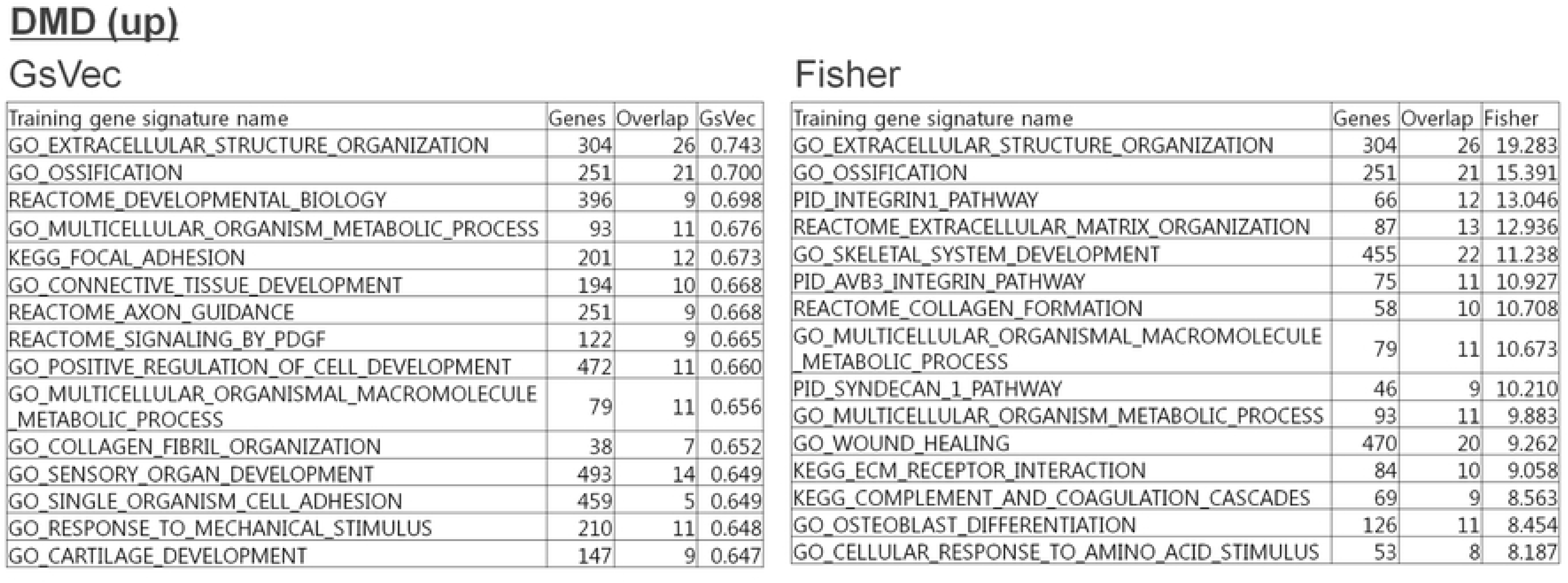
Comparison between GsVec and Fisher of top 15 results with high relevance in DEGs of Duchenne muscular dystrophy (DMD). Using the DEGs of DMD skeletal muscle (E-GEOD-3307) as validation data, each top 15 result that was highly correlated with GsVec or highly significant with Fisher is presented in the left and right tables.

### Systemic lupus erythematosus (SLE) whole blood (E-GSEOD-72509)

A whole blood dataset from SLE patients was analyzed as representative of autoimmune disease. The data were extracted from a heterogeneous population, including those with high and low interferon signature values. However, both GsVec and Fisher fully identified the signatures of immune-related genes with similar results (see Fig 10).

**Fig 10.**
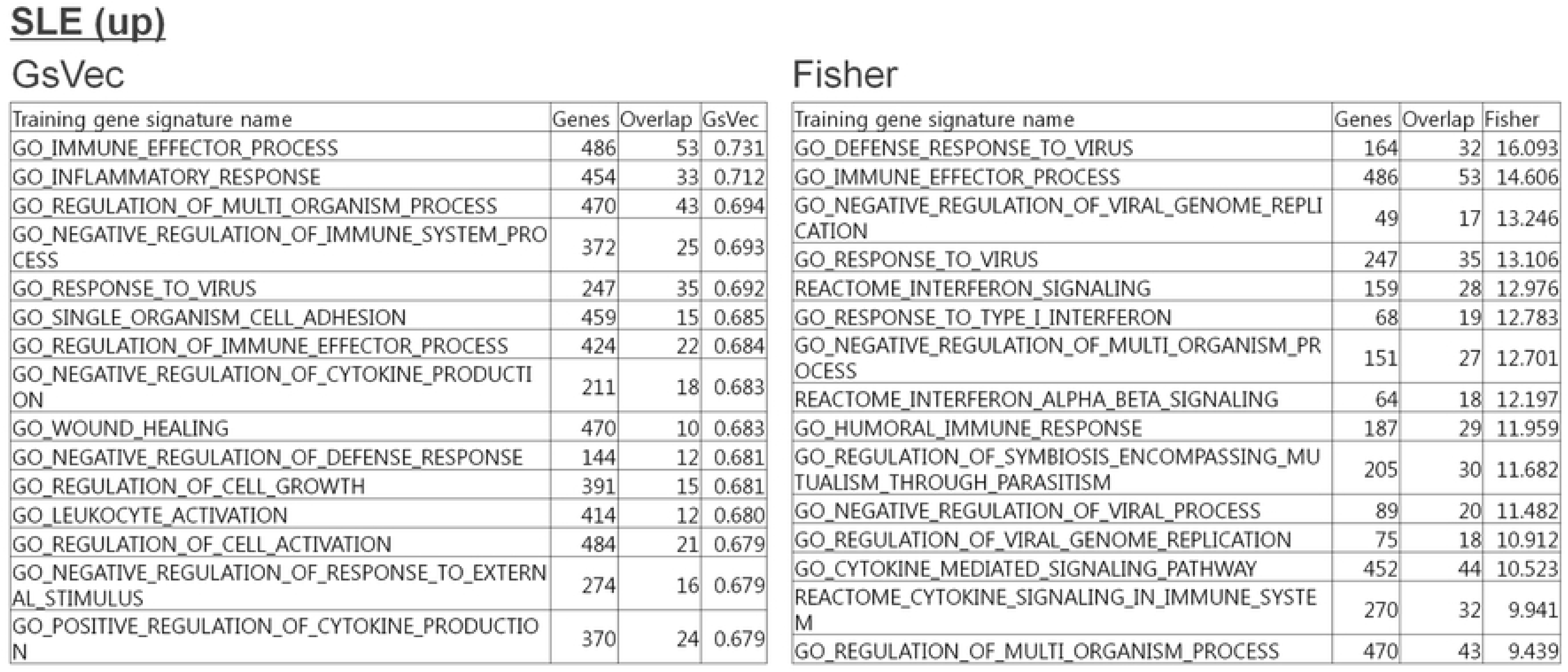
Comparison between GsVec and Fisher of top 15 results with high relevance in DEGs of systemic lupus erythematosus (SLE). Using SLE whole blood (E-GSEOD-72509) DEGs as validation data, each top 15 result that was highly correlated with GsVec or highly significant with Fisher is presented in the left and right tables.

### Sarcoidosis lung tissue (E-GEOD-16538)

We analyzed a dataset derived from lung tissue from sarcoidosis patients. The results indicated that the features of the gene signatures identified by GsVec and Fisher were partially different (Fig 11). In GsVec, gene signatures related to kinase activity, including *MAPK* (mitogen-activated protein kinase), were highly ranked, whereas Fisher identified several gene signatures related to *chemokine*. The original dataset was intended for the investigation of gene regulation of granulomatous sarcoidosis, and it was reported that the gene network associated with the Th1-type response was overexpressed mainly in lung tissue derived from sarcoidosis [30]. In another paper, not only were Th1 cytokines increased in sarcoidosis, but MAPK, especially p38 activation, was found in cells of bronchoalveolar lavage fluid from patients with sarcoidosis [31]. Chemokines were also reported [32]. In summary, sarcoidosis-related gene signatures were identified; however, the two algorithms GsVec and Fisher exhibited different characteristics.

**Fig 11.**
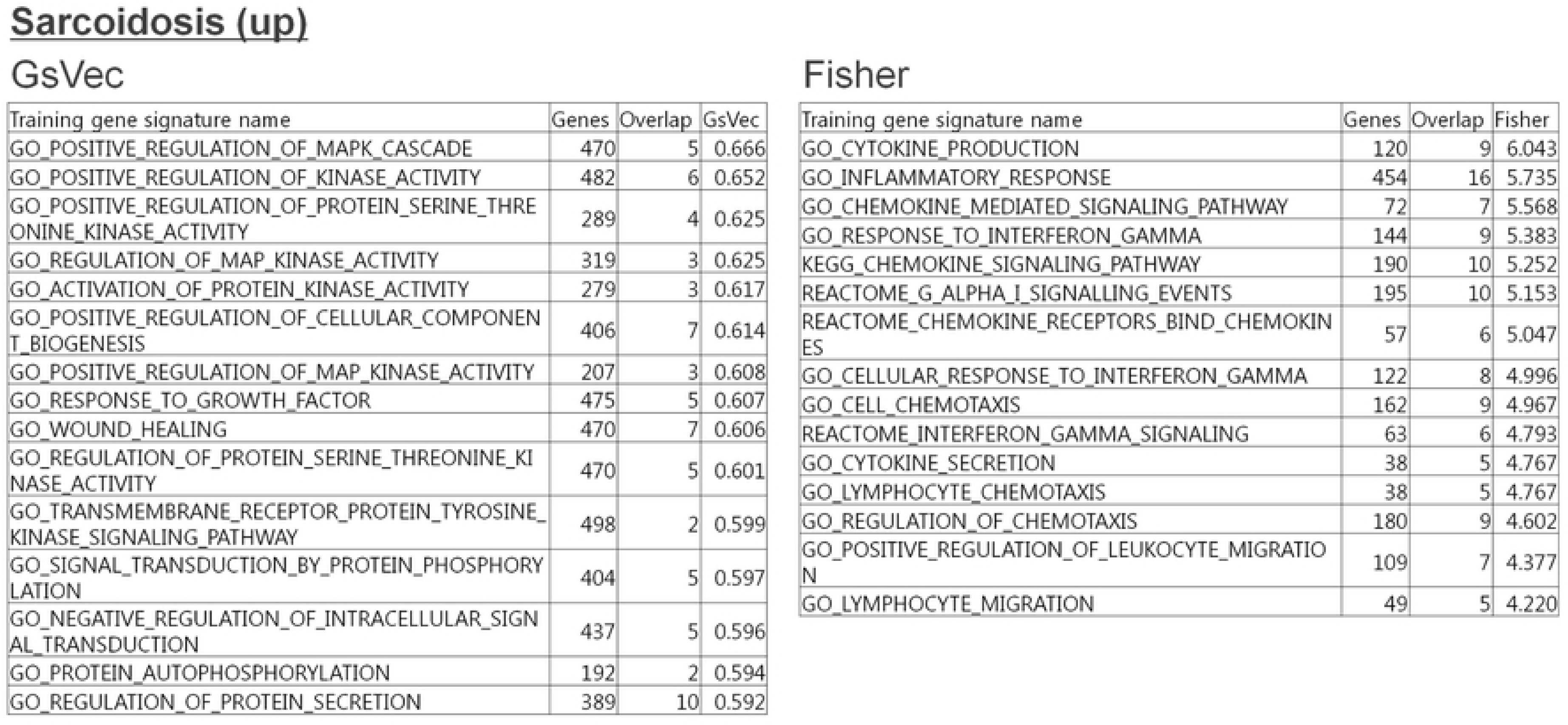
Comparison between GsVec and Fisher of top 15 results with high relevance in DEGs of sarcoidosis. Using the DEGs of sarcoidosis lung tissue (E-GEOD-16538) as validation data, each top 15 result that was highly correlated with GsVec or highly significant with Fisher is presented in the left and right tables.

### Schizophrenia (SCZ) brain Brodmann area 24 (GEOD-78936)

In examining SCZ, we used a dataset derived from postmortem brain tissue. In SCZ GsVec, gene signatures related to *hormone* were highly ranked (Fig 12). *Hormone* was a common but a unique result, which only ranked in GsVec, which suggests the involvement of dopamine in the limbic system [33]. Further, a gene signature, *nervous*, was also highly ranked. In contrast, in Fisher, gene signatures that suggested the involvement of other CNS cells, such as *glia cell* and *astrocyte*, were highly ranked and had different features.

**Fig 12.**
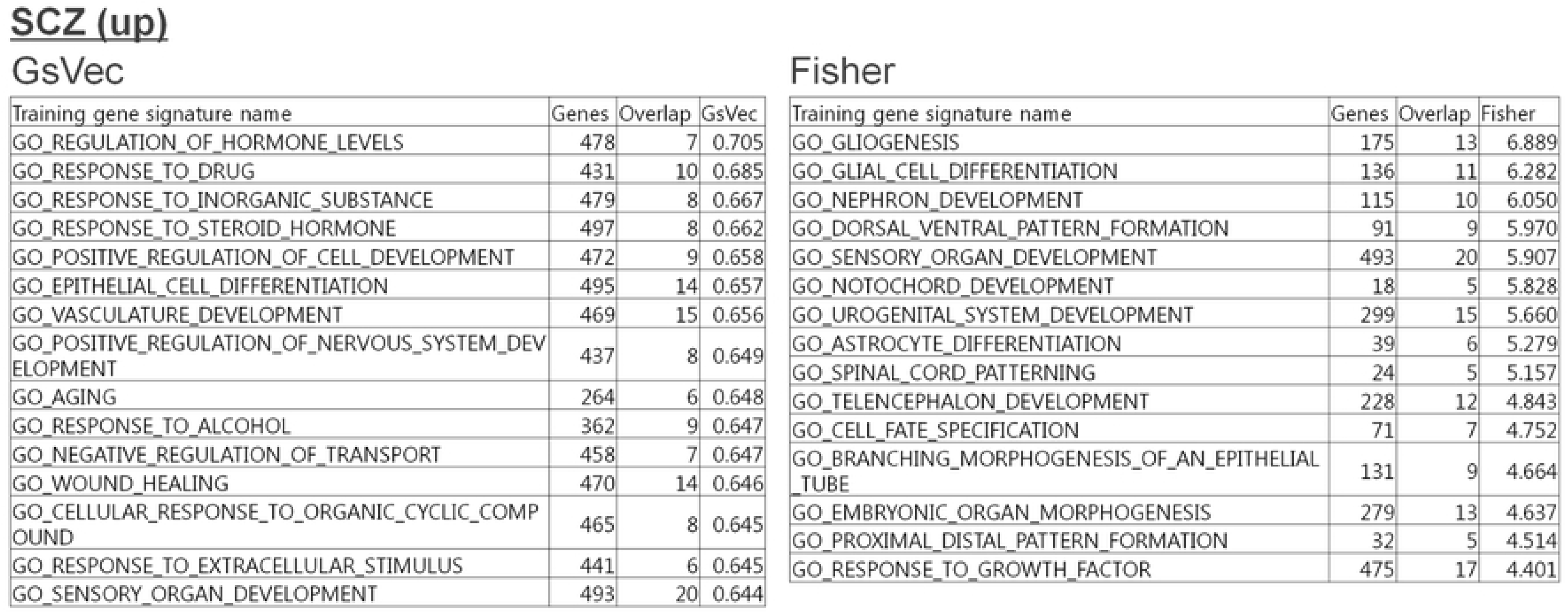
Comparison between GsVec and Fisher of top 15 results with high relevance in DEGs of schizophrenia (SCZ). Using the DEGs of SCZ brain Brodmann area 24 (GEOD-78936) as validation data, each top 15 result that was highly correlated with GsVec or highly significant with Fisher is presented in the left and right tables.

## Discussion

In this paper, we propose a method for associating gene signatures by feature extraction using an NLP method. This method is entirely different from traditional pathway enrichment analysis for gene signature interpretation. Biologically reasonable results were obtained both in the verification of the pathway database (BioCarta) with known biological significance and in the verification using DEGs extracted from the actual gene expression. Compared to conventional pathway enrichment analysis by Fisher’s exact test (Fisher), the proposed algorithm (GsVec) can identify a signature that is equivalent or more biologically relevant than Fisher.

Among diseases known to be related to immunity, GsVec tends to differentiate well between the biological features of autoimmune diseases. In MS, GsVec extracted more biologically valid signatures from immunological and neuronal aspects. Additionally, in CD, the signature called *interferon gamma* and other characteristics (e.g., *cell trafficking*) can be extracted in GsVec. Moreover, in SCZ, a unique signature, *hormone*, was highly ranked only in GsVec. Thus, GsVec captured the signatures related to periphery and CNS more specifically than Fisher.

In this study, there were many reasons for selecting the SCDV-based method among the many methods related to distributed representation of documents in NLP. In the advance analysis, in comparing multiple methods, BOW and TFIDF did not consider gene similarity that was not directly overlapping, resulting in a high correlation with Fisher; thus, the advantage of using NLP methods was low. In the averaging of word2vec (gene vector), the specificity of a gene signature was not taken into account, and thus did not meet our purpose. In addition, as a result of assuming a general bioinformatics analysis environment (e.g., R language, PC specifications) as the potential of this research development, methods with a large amount of computation using deep learning were excluded from the candidates, as well as methods that were difficult to implement in the R language. Based on these considerations, the SCDV method was considered to be an optimal method that could be executed in a general bioinformatics analysis environment while capturing the characteristics of gene signatures.

However, there are several problems with this approach. First, it is difficult to determine whether approximately 5,000 gene signatures are sufficient as training data. In NLP, tens of thousands of data are generally used as training data. However, inadvertently mixing different gene signatures to increase training data (e.g., other non-curation-based collections published in MSigDB) can adversely affect the quality of the signature vector. Further enhancement of pathway data with clear biological meaning is thus necessary.

Second, NLP can identify many words that appear in a specific document as important, but gene signatures do not duplicate genes in signatures; thus, the weight of important genes may be insufficient. This may create a discrepancy with human intuition regarding the key gene in the gene signature (pathway).

The third problem is a general problem in machine learning and artificial intelligence [34]. The relationship between signatures indicated by GsVec has strong elements that cannot be expressed by direct gene duplication; thus, it may be difficult to specify the rationale. Therefore, it may be desirable to combine GsVec with a well-grounded Fisher or other statistical method instead of using it on its own.

Despite the aforementioned problems, the proposed method demonstrates results that are equivalent or superior to those of conventional methods, and has high potential. Training data improvements, feature vectorization and topicalization methods, and identification of important genes are examples of potential improvements.

In the future, if the pathway database is generalized considering the direction of the regulatory relationship of genes, NLP methods that focus on context and learn to sequence from the beginning of sentences can also be applied in this field. Several NLP platforms are already able to graph regulatory relationships between genes (e.g., IBM Watson for Drug Discovery). Improvements in the accuracy of these platforms will increase the value of methods that use NLP and are capable of biological interpretation close to human performance.

## Methods

### Preparation of training and validation data

A total of 5,254 gene signatures were used as a learning set whose genes fell within the range of 10 to 500 genes from KEGG and REACTOME in the C2 canonical pathway and C5 GO biological process sets in MSigDB. Similarly, 214 gene signatures of BioCarta in the C2 curated gene set were used as the validation set. For these signatures, only relevant gene signatures were extracted from the gtm file provided by MSigDB and saved in the same gtm format as MSigDB.

The following operations were performed using the R language integrated environment Microsoft R version 3.5 and R studio version 1.1.463. The created gtm file was converted to data.frame listing each gene signature’s unique ID, name, description, number of genes, and gene symbol, and was output as a text file. The above operations can be executed in one step as the original R functions *make_train.data* (for training data) and *make_validation.data* (for validation data).

### Creation of gene vectors

Gene feature vectors (gene vectors) were created using the R fastText package [35]. The number of characters used for subwording and the number of preceding and following words analyzed as related words was set to 10,000, and gene vectors were created by the co-occurrence of the entire gene signature without using functions for subwording, preceding and following words. First, a 1/0 matrix (one-hot vector) of genes × gene signatures based on the presence/absence of corresponding genes was created. Then, a large 0/1 matrix of combinations of the number of genes and gene signatures was formed. The matrix was compressed to a low dimension by the skip-gram model using negative sampling for the co-occurrence probability of genes that appeared simultaneously in the gene signature. It was then designated as a gene vector. Fig 2 presents *gv_i_*, a vector of an arbitrary gene *g_i_*.

The number of dimensions to be compressed (i.e., gene vector length) and the number of learning iterations (epoch number) had to be adjusted according to the number of vocabularies in the NLP analysis. Because of the advanced analysis, the number of dimensions was set to 150 and the number of epochs to 100 (see S1 and S2 Figs). The above operations can be executed in one step as an original R function, *gs.train_genevec*.

### Creation of gene-topic vectors

Topics (clusters) included in these gene signatures were extracted from the created gene vector by soft clustering using the Gaussian mixture model (GMM). For the GMM analysis, *mclust* of the R mclust package [16] was used, and the number of clusters was estimated using the *mclustBIC* function. The GMM analysis was performed by the *mclust* function with the number of clusters determined by the Bayesian information criterion (BIC), and the probability that each gene contributed to each cluster was calculated. This probability was multiplied for each cluster by the previous gene vector to obtain a gene-cluster vector. Fig 2 demonstrates 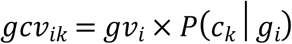, where *c* represents a cluster and *K* represents the number of clusters.

Separately, a score to reduce the weight of genes that appeared in various signatures was calculated by determining the value obtained by dividing the total number of signatures for each gene by the number of signatures that contained the gene from the one-hot vector of the genes × gene signatures (hereinafter referred to as the *inverse signature factor*). The following equation was expressed as a function of R as a countermeasure to infiniteness; when the gene was 0, the weights were normalized.

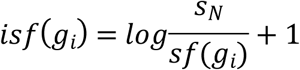

Here, *S_N_* represents the total number of gene signatures, while *Sf* (*g_i_*) represents the number of gene signatures including the gene *g_i_*.

By multiplying this value by the previous gene-cluster vector, gene-topic vectors reflecting the height of gene contribution to each topic and the gene specificity weight were calculated. Fig 2 displays 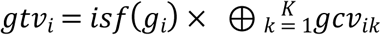, where ⊕ represents concatenation.

In addition, the *estimate_cluster_size* function for estimating the number of clusters from the gene vector and the *gs.train_topicvec* function for creating the gene-cluster and gene-topic vectors were created, making executable in one step. The validation results of the parameters for estimating the number of clusters are presented in Supplemental Fig 3.

### Creation of signature vector

Training data or validation data were input to the generated gene-topic vectors, and the gene-topic vectors were averaged for the genes included in each gene signature. As a result, each signature-specific feature vector (hereinafter referred to as the signature vector) was created, taking into account the gene specificity and the relevance of the gene to the topic in the training data. Fig 2 demonstrates 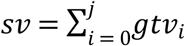. The above operation can be performed with an original function, *predict_GsVec_from.TopicVec*.

It should be noted that the original SCDV method of NLP, which is the basis of this method, can increase the speed and accuracy using the sparse method [9]. However, in gene signature analysis, the number of genes corresponding to the number of vocabularies is overwhelmingly small compared to natural language; thus, this step was excluded because the above procedure neither increased speed nor improved accuracy.

### Association between signature vectors by GsVec

The association between the signature data of the training and validation data was calculated based on the cosine similarity score of the signature vector. Depending on the combination of training and validation data, a large amount of computation is required; thus, the existing cosine distance function of the R package was not used, and high-speed program code was created by original matrix computation.

This operation can be performed with the original functions *similarity_vectors* and *GSVEC*. However, while the former outputs minimal results, the latter is a comprehensive function with various options, such as adding annotations of the original signature and simultaneously outputting the results of Fisher.

### Conventional pathway enrichment analysis by Fisher’s exact test

Fisher’s exact test created and implemented its own function, *gs.enrich_fisher*, to perform comprehensive processing between gene signatures using the *fisher.test* function of the R stats package.

### Visualization of GsVec results with tSNE

To visualize the similarity by the cosine distance between the signature vectors of GsVec, the cosine distance matrix was first linearly compressed with principal component analysis (PCA) using the *prcomp* function of the R stats package. The top 95% of the principal components were projected in two dimensions using the *Rtsne* function in the R Rtsne package [36] and visualized using the R ggplot2 package [37]. This series of operations can be executed with the original function *pca.tsne_GsVec*.

### Extraction of DEGs from public gene expression data

The DEGs of representative diseases were selected from several datasets in which the gene expression of the appropriate disease site was used, and which was a representative disease from various disease areas among the already calculated DEGs published in the Expression Atlas [19]. With regard to cancer, the Expression Atlas did not provide an appropriate dataset; therefore, cancer tissue and matched normal tissue datasets of major cancer types were taken from the TCGA database [20]. TCGA data were normalized from the RNA-seq count data using the *voom* method in the R limma package, and statistically tested by the experimental Bayes method. DEGs with a false discovery rate -adjusted P-value of 0.001 or less and a fold change of ±2 or more were extracted.

### Publishing program codes

The program code for the GsVec analysis developed in this study is freely available from https://github.com/yuumio/GsVec

## Acknowledgments

We would like to show our greatest appreciation to Prof. Masami Hagiya, Dr. Rei Kawakami, and Dr. Toshiaki Nakazawa of the University of Tokyo AI Data Frontier Course who taught us about AI, machine learning, and NLP. We would like to offer special thanks to Dr. Shinichi Kondo and Dr. Shuji Sato for supporting this research. The authors would like to thank Enago (www.enago.jp) for the English language review.

## Supporting information

**S1 Fig.**
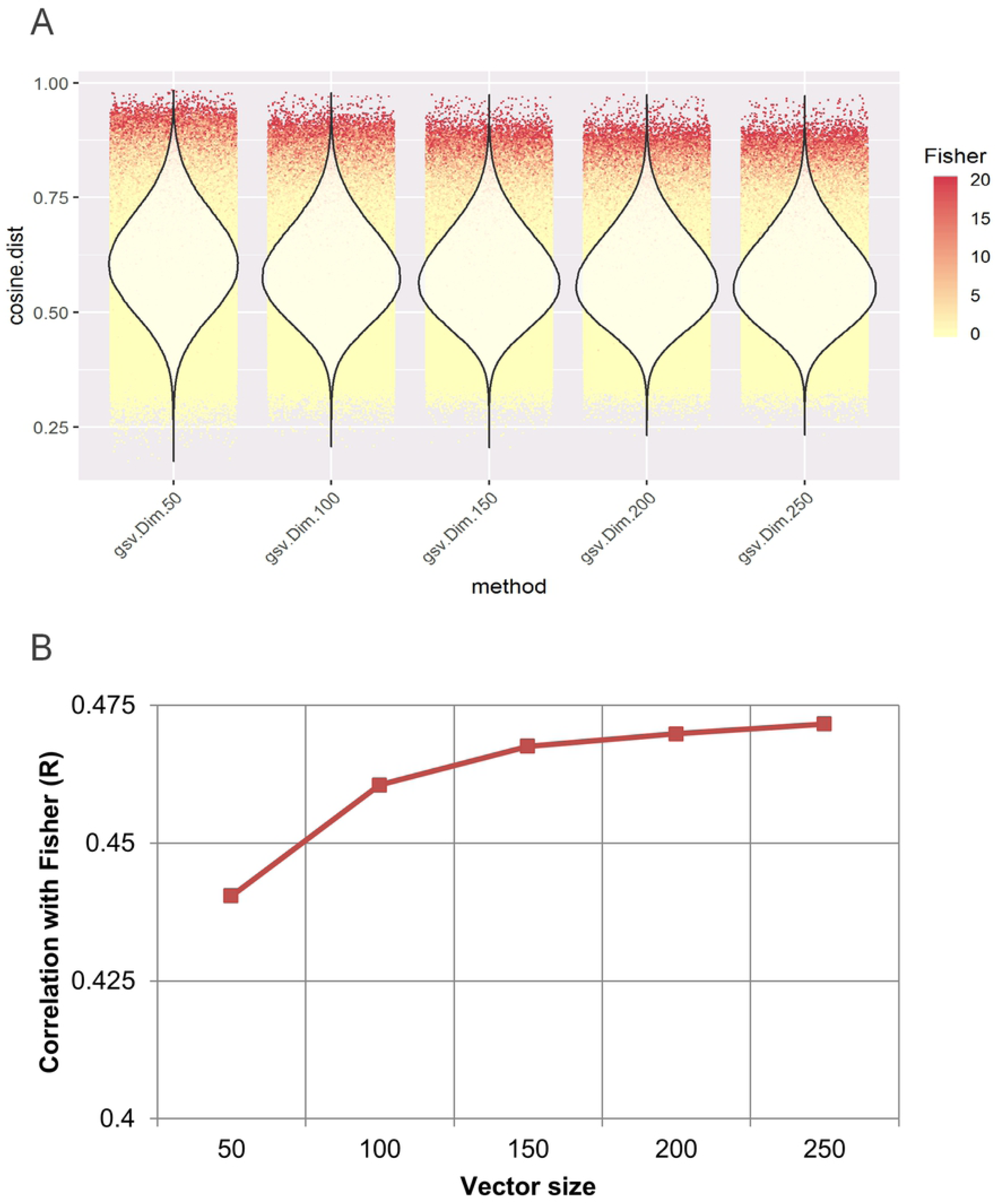
Examination of dimensionality condition in gene vectors. Gene vectors were created for five vector sizes of the gene vectors: 50, 100, 150, 200, and 250. A) Similarity with the validation data was calculated. The distribution is illustrated in a violin plot, while the Fisher result is presented in a scatter plot. B) The Pearson correlation coefficient with Fisher for each vector size is presented as a bar graph.

**S2 Fig.**
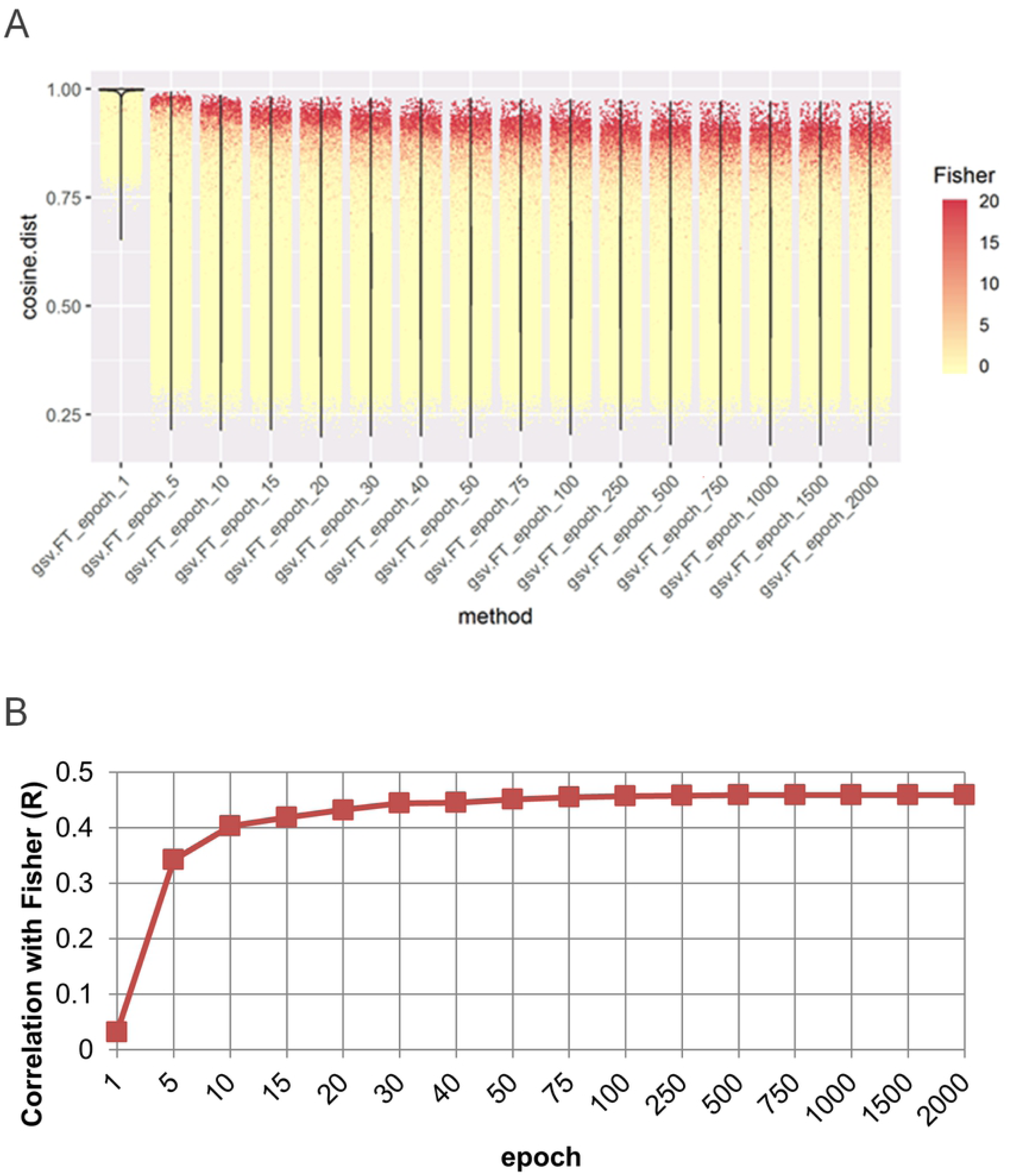
Examination of epoch number in gene vectors. Gene vectors were created for 16 levels: 1, 5, 10, 15, 20, 30, 40, 50, 75, 100, 250, 500, 1,000, 1,500, and 2,000. A) Similarity with the validation data was calculated. The distribution is presented in a violin plot, while the Fisher result is presented in a scatter plot. B) The Pearson correlation coefficient with Fisher for each vector size is illustrated as a bar graph.

**S3 Fig.**
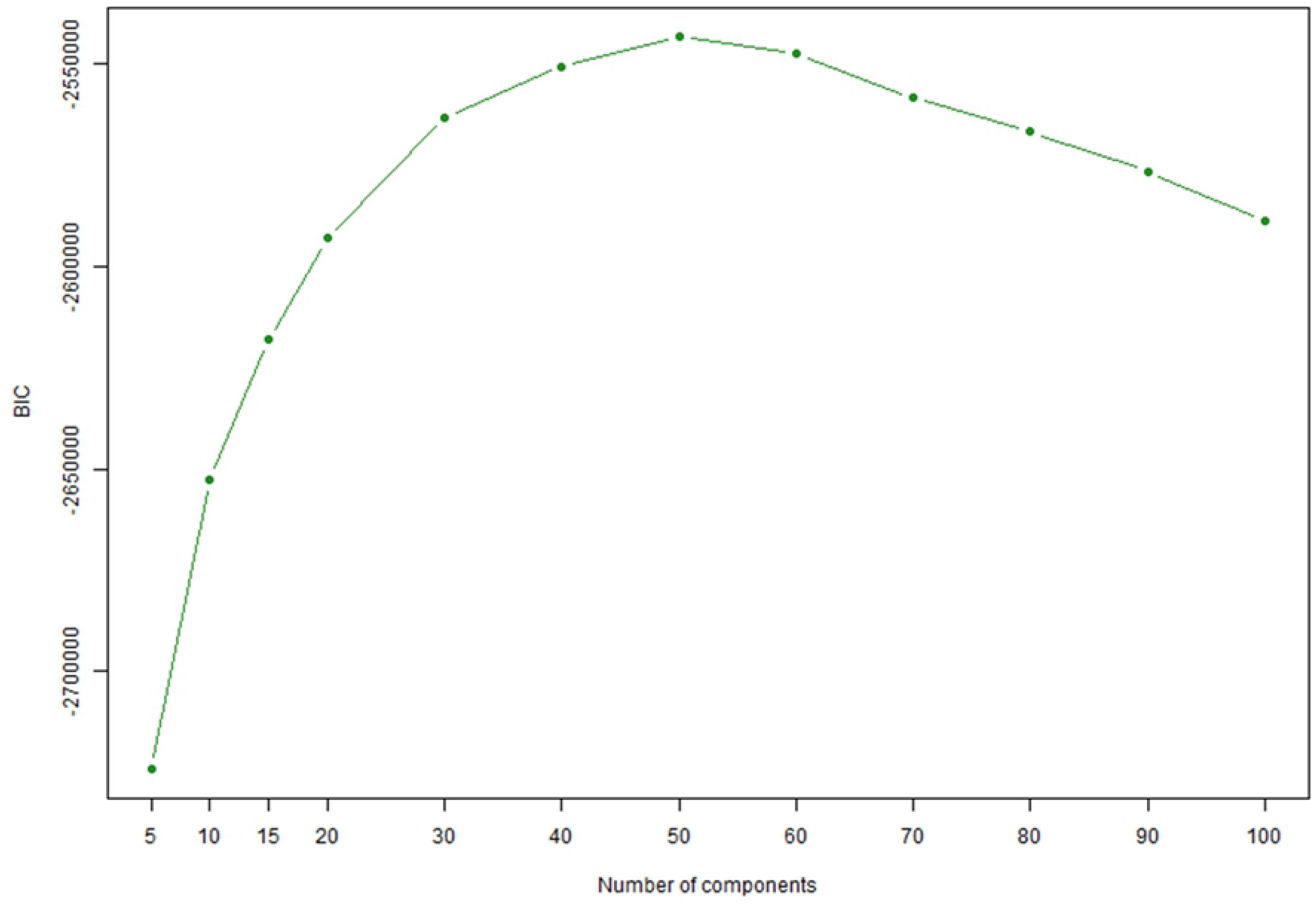
Validation of parameters for estimating the number of clusters as topics in gene signatures. The output of the *mclustBIC* function of the R mclust package was visualized by the *plot* function. The Bayesian information criterion in the mclust package is 2 × log likelihood. Thus, the largest value was selected as the optimal cluster.

